# The ecology of carbon sequestration in coastal upwelling ecosystems

**DOI:** 10.64898/2025.12.29.696936

**Authors:** Michael R. Stukel, Lihini I. Aluwihare, Daniele Bianchi, Heather J. Forrer, Monique Messié

## Abstract

The biological carbon pump (BCP) transports carbon fixed by phytoplankton into the deep ocean via multiple pathways and leads to net carbon dioxide sequestration. The BCP is very active in highly productive eastern boundary upwelling systems (EBUSs), however, the extent to which climate change will impact this key ecosystem service remains uncertain. While this review and synthesis focuses on results from the California Current Ecosystem (the most extensively studied EBUS), similarities and differences are also noted with the Benguela, Canary, and Humboldt ecosystems. We focus on vertical carbon export mediated by sinking particles (responsible for >half of the BCP in EBUSs), as well as subduction of organic matter and vertically migrating zooplankton and fish. We suggest that a plug-flow-reactor conceptual model can be used to link BCP results with physical circulation changes within EBUSs to predict whether climate change will lead to increased or decreased biologically mediated CO_2_ uptake.

## Background

### Primary production, new production, and the ocean’s biological carbon pump

The deep ocean is a vast reservoir for dissolved inorganic carbon (CO_2_, H_2_CO_3_, HCO_3_^-^ and CO_3_^2-^). It is enriched in inorganic carbon relative to the surface ocean as a result of the solubility and biological carbon pumps (Volk and Hoffert 1985). The solubility pump is the downward transport of inorganic carbon driven by the simultaneous high solubility of CO_2_ in cold water and the high density of cold water, which leads it to sink into the ocean interior. The biological carbon pump (BCP, Fig. 1) refers to the downward transport of organic matter as the result of complex interactions between ocean circulation, biogeochemistry, and ecology (Longhurst and Harrison 1989, Ducklow et al. 2001, Boyd et al. 2019). The BCP begins with the conversion of CO_2_ into organic matter by photosynthesis in phytoplankton (floating algae) in the ocean’s euphotic zone (the sunlit surface layer that typically varies from 20 – 150 m thick). While most of the carbon fixed by phytoplankton will be respired in the surface ocean, a portion of it escapes respiration and is transported into the deep ocean within sinking particles (Martin et al. 1987, Buesseler and Boyd 2009), through the diel and seasonal vertical migrations of zooplankton and micronekton (Steinberg et al. 2000, Davison et al. 2013, Jónasdóttir et al. 2015), and by passive transport of suspended particulate and dissolved organic matter during subduction and vertical mixing (Carlson et al. 1994, Levy et al. 2013, Omand et al. 2015, Lacour et al. 2023). Together these processes are estimated to transport between 5 and 13 Pg C yr^-1^ out of the euphotic zone globally (Laws et al. 2011, Archibald et al. 2019, Henson et al. 2022, Nowicki et al. 2022).

**Fig. 1.**
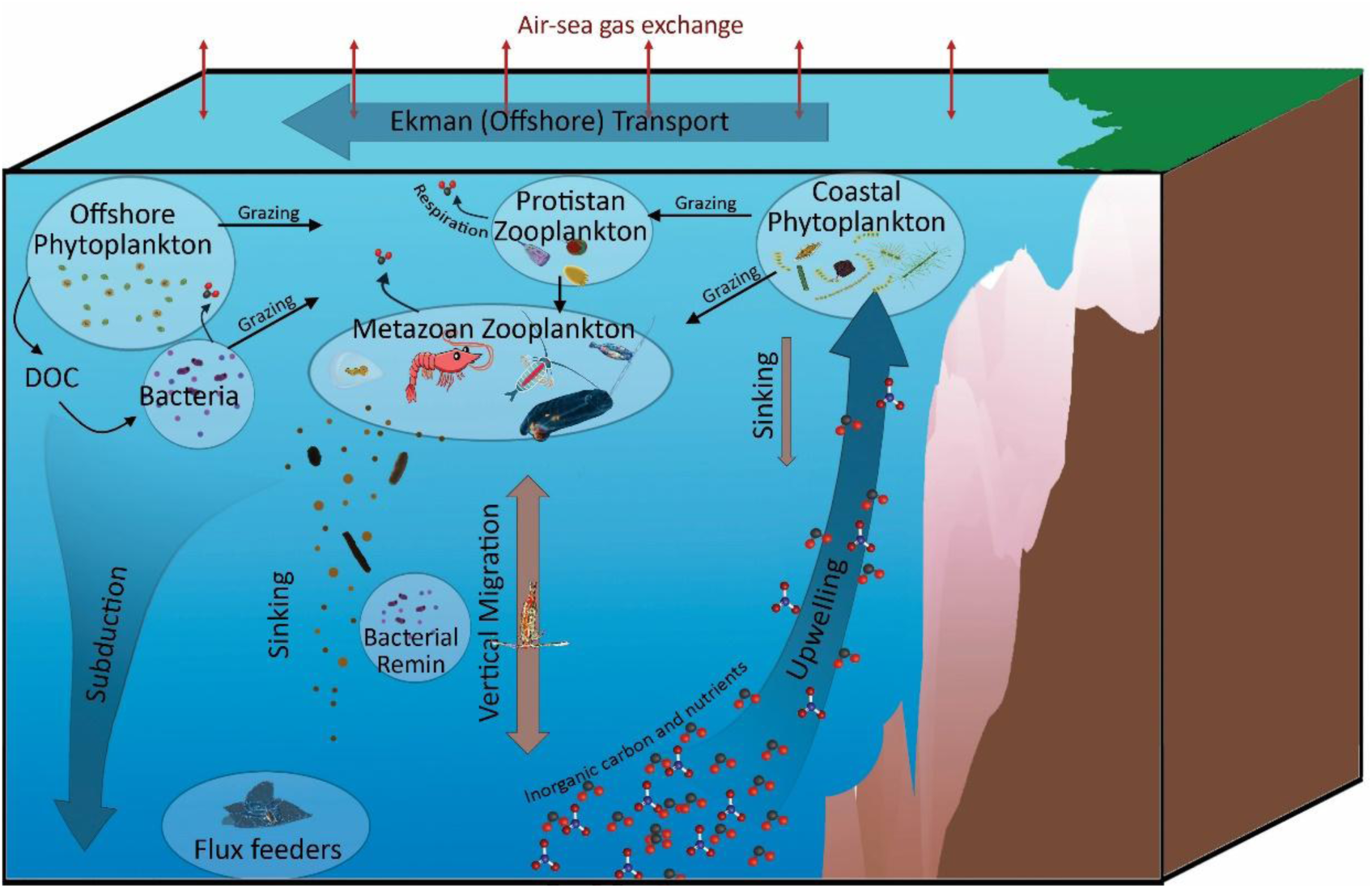
The biological carbon pump (BCP) in an eastern boundary current upwelling ecosystem. Wind-driven upwelling brings nitrate and inorganic carbon (including CO_2_) into the surface ocean. The upwelled nitrate stimulates phytoplankton production (“new production”) near the coast. This coastal phytoplankton community is typically dominated by large taxa including diatoms that can contribute to direct sinking either as individual cells or within aggregates. The plankton community evolves as it is transported offshore by Ekman transport. Protistan and metazoan zooplankton both graze on the phytoplankton community, with the latter primarily serving to regenerate nutrients (e.g., ammonium) and respire carbon dioxide. The metazoan zooplankton community (and higher trophic levels), in contrast, can contribute substantially to the BCP both through the production of rapidly sinking fecal pellets and through active transport driven by their vertical migrations and respiration in the mesopelagic realm. As the community continues to evolve and nutrients become exhausted, smaller phytoplankton tend to dominate. These taxa are less likely to sink on their own and are predominantly grazed by protistan zooplankton, presumably leading to a less efficient BCP. All phytoplankton and zooplankton also produce dissolved organic carbon (DOC) through multiple processes that stimulate bacterial production and respiration as part of the “microbial loop”. Sinking carbon flux typically declines with depth due to both the respiration of particle-attached microbes and zooplankton detritivory (including the impacts of flux-feeding zooplankton). Another important aspect of the BCP in coastal upwelling biomes is subduction, which transports entire water parcels (including organic carbon in living organisms, detritus, and dissolved molecules) into the deep ocean. Subduction can be driven by meso- and submesoscale processes (e.g., fronts and eddies) or by the large-scale ocean circulation.

Because marine organic matter has a relatively narrow carbon-to-nitrogen (C:N) stoichiometric ratio (106 mol C: 16 mol N, Redfield 1934), the downward flux of organic matter, referred to as ‘export production’, must be balanced by an equivalent supply of ‘new’ nutrients, referred to as ‘new production’. This nutrient supply primarily occurs through upwelling of nutrient-rich water, dissolved N_2_ gas fixation by diazotrophs, atmospheric deposition, and coastal inputs including rivers, groundwater, and anthropogenic sources (Dugdale and Goering 1967, Eppley and Peterson 1979, Doney et al. 2007, Kessouri et al. 2021). While this mass balance between export and new production must hold when integrating over sufficiently large spatial and long temporal scales, local imbalances occur at daily to seasonal time scales due to time lags associated with the sequence of production. Specifically, delays between 1) nutrient input, 2) phytoplankton growth, 3) aggregation of phytoplankton into marine snow or zooplankton grazing and subsequent fecal pellet production, and 4) sinking of organic matter create a temporal decoupling of new and export production. This presents challenges when attempting to close BCP budgets, which may be further complicated by spatial decoupling of these processes within advective ecosystems (Plattner et al. 2005, Stukel et al. 2015a, Messié et al. 2025).

Global climate models vary in their predictions of both the current magnitude of and future changes in the BCP, although most project a future decline in sinking particle flux (Henson et al. 2022). When assessing the role of the BCP as a feedback in the global climate system, however, a focus on the magnitude of export flux is insufficient (Frenger et al. 2024). Indeed, most climate models predict that increased stratification – and the associated reduction in supply of new nutrients – is a dominant driver of the projected decline in BCP magnitude. Yet, reduced upwelling introduces less CO_2_ into the joint surface ocean-atmosphere system, potentially offsetting the decline in export production. The BCP will thus only act as a feedback on climate change if the stock of biologically derived carbon in the deep ocean changes (Frenger et al. 2024). Importantly, changes in the stock will not necessarily be predicted directly by changes in the overall magnitude of the BCP, but also by changes in the C:N (or carbon:phosphorus, C:P) ratios of exported organic matter relative to upwelled dissolved molecules (i.e., inorganic carbon and nutrients). Such changes could be mediated by 1) nitrogen fixation or denitrification (each of which modify the bioavailable nitrogen pool, Zehr and Kudela 2011, Casciotti 2016), 2) variability in pre-formed nutrient concentrations in subducted waters (i.e., nutrients that are not utilized before water masses sink, often as a result of trace element limitation, Martin 1990), 3) modified elemental stoichiometry of sinking particles (which can occur either as a result of different particle origin or because of differential utilization of elements by particle-degrading microbes, Knauer et al. 1979, Tanioka et al. 2021, Lomas et al. 2022), or 4) changes in the relative importance of different BCP pathways which might have different stoichiometries (for instance dissolved organic matter transported by the subduction pump may have much higher carbon:nitrogen than sinking particles, Aminot and Kérouel 2004, Letscher et al. 2015). It is thus crucial to assess how multiple BCP pathways may be changing in major marine biomes in response to shifting climatic conditions (Boyd 2015, Henson et al. 2024).

#### Box 1

##### Ocean carbon pump terminology

**Solubility carbon pump** – Net downward flux of inorganic carbon (i.e., carbon mass per unit area per unit time across a horizontal surface) mediated by the ocean’s overturning circulation. Because CO_2_ is more soluble in cold water and cold water is dense, waters that sink into the ocean’s interior are typically enriched in CO_2_.

**Biological carbon pump (BCP)** – Net downward flux of organic carbon originally fixed by phytoplankton that enters the ocean interior either within sinking particles, through the vertical migrations of organisms, or when water sinks into the ocean’s interior carrying organic matter with it.

**Gravitational pump** – Downward carbon flux associated with sinking particles and aggregates including phytodetritus (dead phytoplankton), fecal pellets of zooplankton and fish, and carcasses. Particles can sink at rates varying from m d^-1^ to km d^-1^, with sinking speeds controlled by density (especially mineral ballasting), shape, and size (large particles sink faster).

**Active transport** – The portion of the BCP mediated by the vertical migrations of swimming organisms (primarily zooplankton and fish). Active transport is predominantly driven by diel vertical migrators that feed in surface layers during the night and descend to depth to avoid visual predators during the day. It also includes ontogenetic vertical migrators (the “lipid pump”) that migrate seasonally as a part of their life cycles.

**Subduction pump** – The portion of the BCP driven by downward water movement that carries organic matter (both dissolved and particulate, living and dead) into the deep ocean. Subduction can be driven by large-scale circulation features, such as downwelling that occurs across the broad oligotrophic ocean gyres, or at meso- and submesoscale fronts and eddies. Downward organic matter transport is also driven by deep convective mixing (i.e., deep winter mixed layers).

**Sequestration timescales** – The length of time that exported carbon is stored within the ocean can vary from less than a year to millennia and is strongly correlated with the depth to which the carbon is exported.

**New production** – Biological production supported by nutrient sources that are external to the sunlit surface ocean. In the CCE upwelled nitrate is the primary “new” nutrient, although nitrogen fixation and coastal nutrient inputs are also new nutrient sources. Mass balance requires that over sufficiently large spatial and long temporal scales, new production must balance the amount of nitrogen exported as part of the BCP. Since C:N elemental ratios of marine organic matter are fairly well constrained, new production thus puts constraints on the total carbon exported by the BCP.

**Euphotic zone** – Sunlit surface ocean region, within which phytoplankton grow and CO_2_ can exchange with the atmosphere

**Mesopelagic zone** – Also sometimes referred to the twilight zone, this is a mid-depth region in the ocean (200 – 1000 m) that is too dim for photosynthesis and within which substantial carbon flux attenuation occurs.

### Coastal upwelling biomes and the biological carbon pump

Coastal upwelling ecosystems, also referred to as eastern boundary upwelling systems (EBUSs, Fig. 2), are highly active carbon reservoirs with large upward inorganic and downward organic carbon fluxes (Hales et al. 2005, Chavez and Messié 2009, Fréon et al. 2009). Wind-driven upwelling brings CO_2_-rich waters to the surface ocean, potentially leading to net outgassing. CO_2_ upwelling is offset, however, by high photosynthesis rates in surface waters, supported by nutrient input from the same upwelled waters (Ducklow and McCallister 2004, Messié and Chavez 2015). This balance varies with distance from shore, reflecting the time lag between upwelling and productivity, with typical CO_2_ supersaturation and outgassing nearshore and undersaturation and ingassing further offshore (Landschützer et al. 2020, Damien et al. 2023). Long-term CO_2_ storage in deep waters is mediated by the solubility pump and the BCP. The BCP is especially active in upwelling systems, where upwelling introduces nutrient-rich water into the surface ocean and stimulates the production of large phytoplankton taxa, such as coastal diatoms (Bakun 1975, Venrick 2002, Chavez and Messié 2009, Goericke 2011, Taylor and Landry 2018). These phytoplankton communities in turn support abundant stocks of higher trophic-level organisms such as metazoan zooplankton and forage fish, all of which contribute to the BCP through fecal pellet production and diel vertical migrations (Davison et al. 2013, Stukel et al. 2013, Saba et al. 2021).

**Fig. 2.**
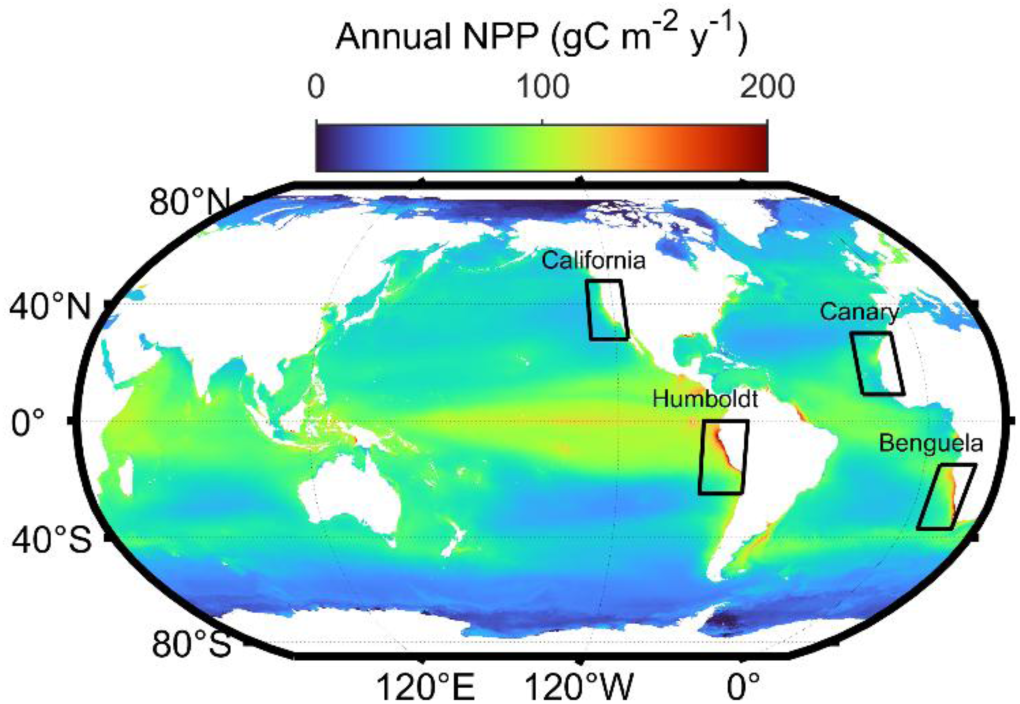
Map of global eastern boundary upwelling ecosystems. Background is satellite-observed annual net primary production from the CAFÉ model (Silsbe et al. 2016).

EBUSs are complex and heterogeneous regions that encompass very strong spatial gradients. While the coastal upwelling domains are some of the most productive pelagic ecosystems in the world ocean, the offshore portions of EBUSs (>200 km from shore) are often strongly stratified and nutrient depleted (Carr and Kearns 2003, Chavez and Messié 2009). There are some distinct differences between the four major EBUSs (Canary, Benguela, Humboldt, and California), including width of continental shelves (narrow in California and Humboldt, wide in Canary and Benguela), latitudinal ranges and seasonality, differences in source waters for upwelling (generally higher nutrient and lower oxygen in Humboldt), and sensitivity to remote interannual forcings (e.g., El Niño-Southern Oscillation). Nevertheless, they all exhibit distinctly similar physical drivers and ecological and biogeochemical responses. Seasonally varying alongshore winds - resulting from the trade winds, the Coriolis force, and land-sea temperature gradients – drive offshore (“Ekman”) transport of surface waters. These surface waters are in turn replaced by upwelled nutrient-rich, CO_2_-rich, and O_2_-depleted deep waters, stimulating high productivity across trophic levels. All EBUSs also feature predominantly equatorward major currents that can displace plankton communities relative to their typical geographical ranges.

In this review, we focus on the California Current Ecosystem (CCE), which is the most well-studied of the EBUSs, while noting that dynamics in this region are broadly similar to those in other EBUSs. The CCE is a natural laboratory for studying the relationships between climate-change, ecology, and biogeochemistry for multiple reasons (Ohman et al. 2013a). It is the site of one of the longest running (75+-years) spatially resolved ocean time-series (California Cooperative Oceanic Fisheries Investigations, CalCOFI, Bograd et al. 2003), which has collected samples spanning ocean physics to zooplankton at stations along the coast and to a distance of ∼500 km from shore from 1949 to the present. It is also the site of the 20+-year CCE Long-Term Ecological Research Program (CCE LTER, Ohman et al. 2013b, Goericke and Ohman 2015), which has added substantial autonomous, remote, and shipboard time-series along with process-oriented analyses of ecosystem dynamics relevant to the BCP. Together, these long-term programs offer an unparalleled record of natural variability and enable detection of climate-driven changes in the physical, biological and chemical dynamics of the CCE EBUS.

The southern CCE (CalCOFI study site, which extends along the coast from ∼San Diego to San Luis Obispo and extends to ∼500 km offshore) encompasses a productive coastal upwelling zone, a transitional zone with moderate and variable productivity, and an oligotrophic offshore domain that is contiguous with the great oligotrophic North Pacific Subtropical Gyre (Venrick 2002, Ohman et al. 2013a, Taylor et al. 2015). Phytoplankton growth and net primary production are predominantly nitrogen limited, although light limitation can occur during coastal upwelling events, and iron (Fe) limitation can arise in the transition zone and at chlorophyll maximum depths (Collier and Palenik 2003, King and Barbeau 2007, Hogle et al. 2018).

Protistan zooplankton are typically the major grazers of phytoplankton (Landry et al. 2009, Connell et al. 2017, Landry et al. 2023); however, a diverse suite of metazoan zooplankton including copepods, euphausiids, and pelagic tunicates are also important (Lavaniegos and Ohman 2007, Ohman et al. 2012, Morrow et al. 2018). These nutrient and ecological dynamics – coupled with physical forcing across multiple timescales up to decadal and long-term secular change (Bond et al. 2003, Bograd et al. 2009, García-Reyes et al. 2014, Jacox et al. 2015, Bograd et al. 2023, Kahru et al. 2023) – shape CCE ecosystem variability. These climatic drivers are linked to long-term changes in the abundances of many zooplankton and forage fish populations, which sometimes manifest as abrupt transitions in the ecosystem (Lavaniegos and Ohman 2007, Rykaczewski and Checkley 2008, Di Lorenzo Emanuele and Ohman 2013, Lindegren et al. 2016, Miller et al. 2019), which likely alter the functioning of the BCP.

Extensive research on the BCP in the CCE has allowed us to understand some of the broad dynamics of carbon export in the region. Early research, for instance, elucidated general patterns in flux attenuation with depth (Martin et al. 1987) and demonstrated seasonal variability in phytoplankton community contributions (variability in pigment and silica) to flux (Landry et al. 1992). More recent studies have paired extensive sediment trap and ^238^U-^234^Th deficiency measurements of sinking particle flux with net primary productivity measurements to assess patterns in efficiency of the BCP (Kelly et al. 2018, Stukel and Barbeau 2020). These results, paired with satellite-observed estimates of net primary production, have identified gravitational sinking to be the most efficient pathway for carbon sequestration in the region, while subduction and active transport were estimated to be weaker components of the BCP (Fig. 3; Stukel et al. (2023)). Responsible for transporting 9.0 mmol C m^-2^ d^-1^ across the 100-m depth horizon in the southern CCE, the gravitational pump sequesters 3.9 Pg C within the 1.7×10^5^ km^2^ study region. In comparison, subduction and vertical mixing were estimated to transport 3.8 mmol C m^-2^ d^-1^ out of the euphotic zone. However, only 0.8 Pg C is sequestered through the subduction pump since the CCE is not a site of deep convective mixing and subducted organic matter is typically remineralized at shallow depths, resulting in shorter mean sequestration times. Active transport by vertically migrating zooplankton and nekton, in contrast, leads to carbon dioxide respired at depths ranging from 200 – 500 m depth (Davison et al. 2013, Ohman and Romagnan 2016, Gastauer et al. 2021). Thus, despite playing a slightly smaller role in export flux from the euphotic zone than subduction (2.9 mmol C m^-2^ d^-1^), it is responsible for slightly more sequestered carbon (1.0 Pg C, Fig. 3, Stukel et al. 2023). Notably, the regional estimates of the subduction pump and active transport are likely underestimates due to the subduction estimates not accounting for the dissolved organic carbon pool, while the active transport estimates omit the contribution of diel vertical migrants that die in the mesopelagic (Stukel et al. 2023). The efficiency of these pathways may be enhanced in other EBUSs, particularly in colder regions with deep convective mixing that drive enhanced subduction through the “seasonal mixed layer pump” (Dall’Olmo et al. 2016) or in regions, such as the Canary Current that may have stronger spring-time subduction related to eddies and/or wind forcing. Additionally, ecosystems containing ontogenetic vertical migrators – species that undergo deep winter hibernation – will have additional contributors to active transport through the “seasonal lipid pump” where deep CO_2_ respiration occurs from stored lipids (Jónasdóttir et al. 2015).

**Fig. 3.**
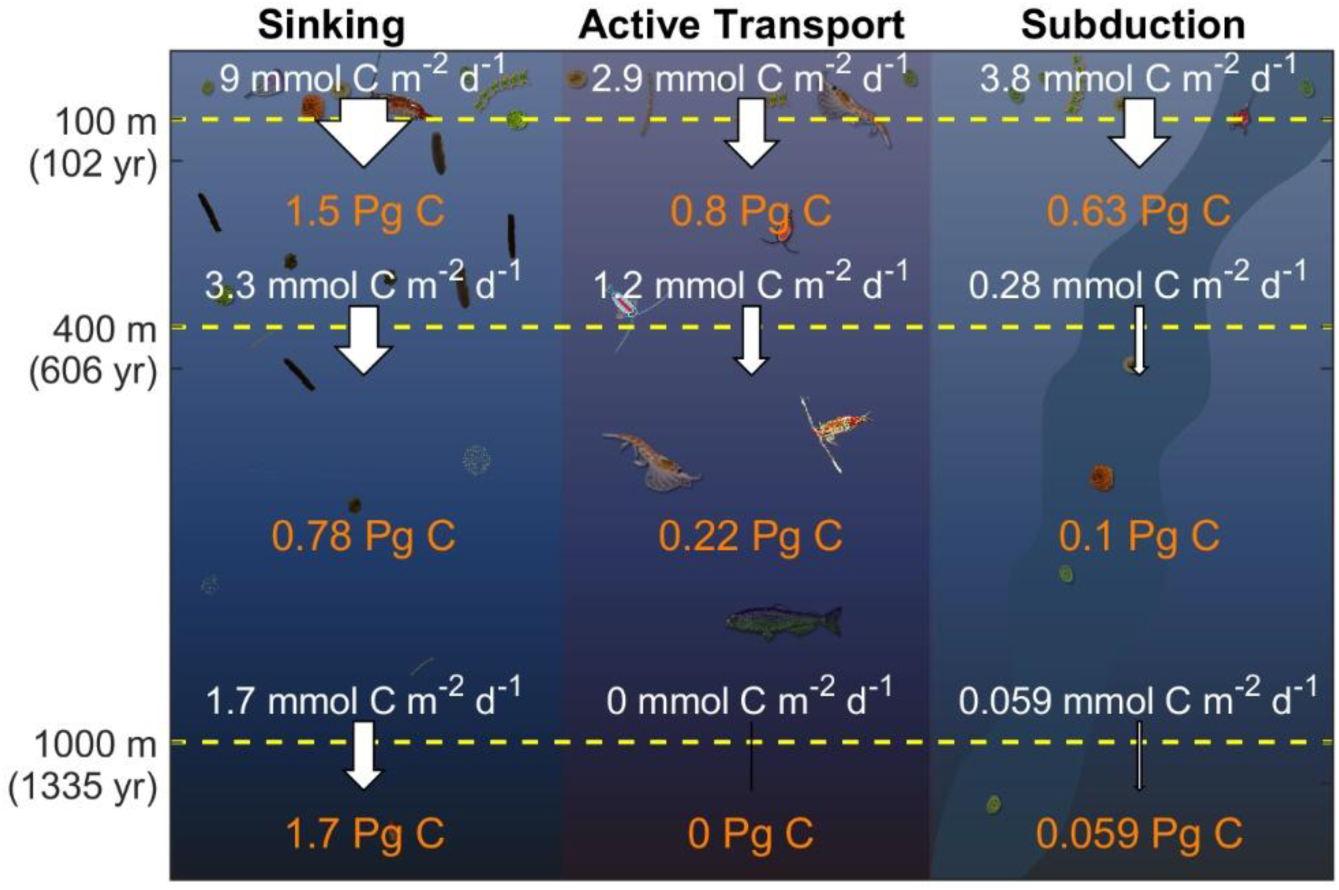
Contributions of gravitational flux (sinking particles), vertically migrating zooplankton and fish (active transport), and passive flux (subduction pump) to export (white arrows and white text) and carbon sequestration (orange text) in three depth horizons in the California Current Ecosystem (modified from Stukel et al. 2023). Note that sinking is the dominant contributor across all depth horizons, while subduction is more important to export across the 100-m depth horizon than active transport, but active transport is more important to carbon sequestration than subduction as a result of deeper remineralization depths.

While the current regional magnitude of the gravitational pump is reasonably well constrained due to extensive, robust quantification and agreement of sinking particle flux measurements from sediment traps and ^238^U-^234^Th disequilibrium (Stukel et al. 2015b, Stukel et al. 2019a), there are some distinct uncertainties about how the gravitational pump responds to changing system productivity. One way to assess these temporal patterns is to focus on predicted changes of new production (uptake of “new” nitrate). These suggest that the *f*-ratio (the ratio of new production to total production where an *f*-ratio ≥0.5 indicates dominant reliance on “new” nitrate; Eppley & Peterson, 1979) is positively correlated with primary production (Fig. 4a, Harrison et al. 1987, Kranz et al. 2020). Such relationships between the *f*-ratio and primary productivity are well supported by basic understanding of phytoplankton ecology. In most ocean regions, periods of high nitrate concentration will correlate to both high total production (often by diatoms) and to a relatively large fraction of that portion being supported by nitrate rather than recycled ammonium. This would suggest that periods of increased upwelling would lead to a more efficient BCP, since export production must balance new production. However, a disconnect is evident between system productivity metrics and export efficiency across EBUSs (Fig. 4b). Specifically, the coastal upwelling biome typically has low export efficiency (5 – 20%), while the offshore oligotrophic domain often has export efficiency >20%. This apparent disconnect between *f*-ratio and export efficiency might be resolved by considering the temporal lags relating new production, horizontal transport, and export (Henson et al. 2015, Laws and Maiti 2019). Pulses of coastal upwelling initiate high rates of new production that typically occur in a relatively narrow band along the coastal region. As the phytoplankton bloom develops, often with subsequent changes in phytoplankton physiological status or blooms of zooplankton, Ekman transport advects the surface water offshore where export production – often reaching peak magnitude during the bloom decline – typically occurs. This introduces a series of lag times (days or even weeks) into the system that can obscure patterns derived from contemporaneous measurements of new and export production. Estimates of these lag times in the CCE can be quite variable and likely relate to temperature and ecosystem structure, ranging from approximately one week to one month (Messié and Chavez 2017, Kahru et al. 2020, Stukel et al. this issue). This spatiotemporal decoupling of new and export production has been predicted by numerical models (Olivieri and Chavez 2000, Plattner et al. 2005, Frischknecht et al. 2018, Messié et al. 2025) and observed in the CCE region (Stukel et al. 2011, Kranz et al. 2020, Chabert et al. 2021).

**Fig. 4.**
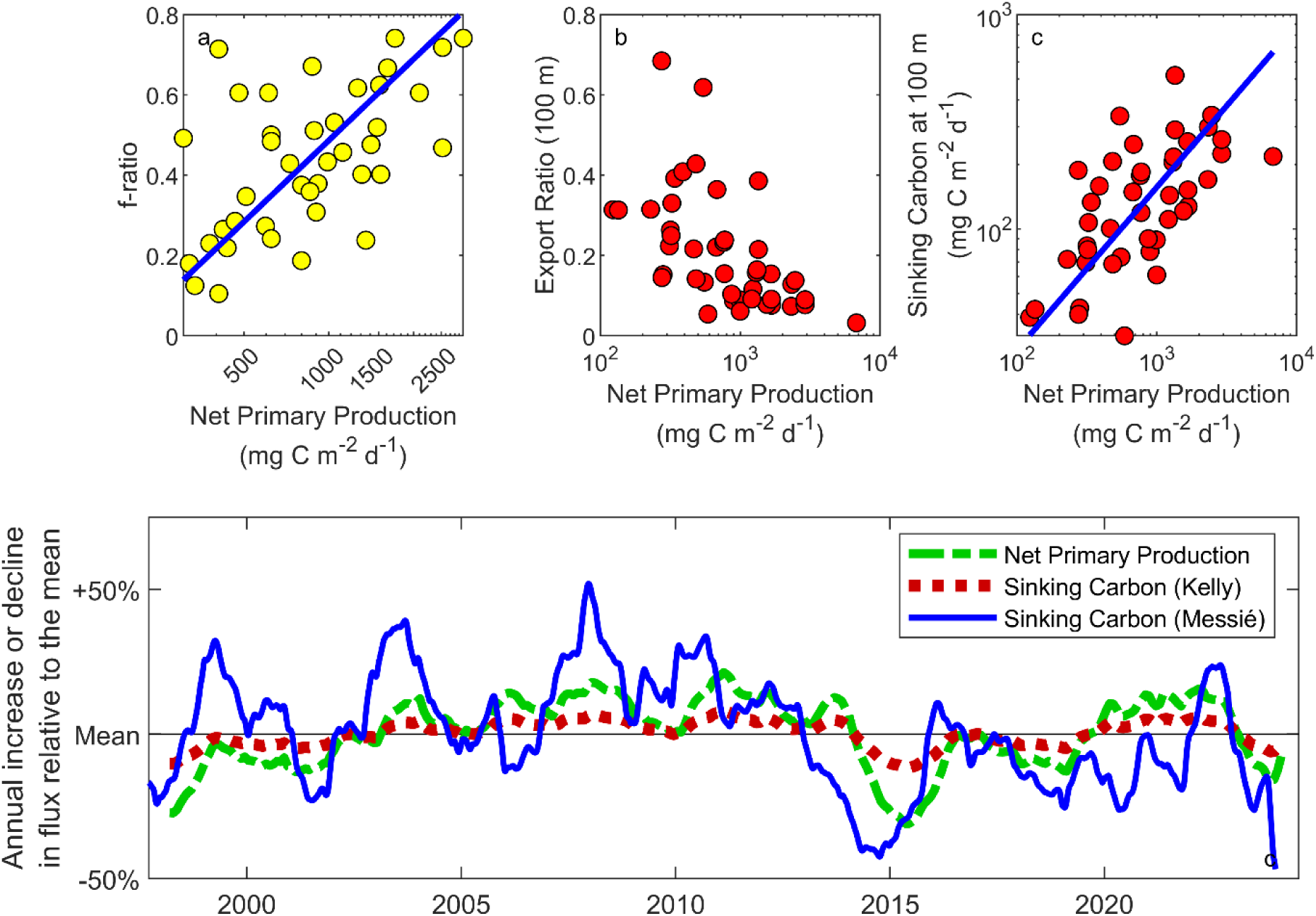
Export efficiency and predicted interannual variability in export flux. a) f-ratio = new production divided by total production as a function of net primary production (NPP) (data from Eppley et al. 1979). Regression line is *f* = *m*×NPP + *b*, where *m* = 0.67 ± 0.1 and *b* = −1.5 ± 0.3. b) Export ratio = sinking particle export (100-m depth horizon) divided by NPP plotted as a function of net primary production (data from CCE LTER sediment traps with methods described in Stukel et al. 2015b, Stukel and Barbeau 2020). Note that we do not plot a regression in panel b, because the export ratio is correlated with net primary production. c) Export as a function of NPP. Regression line is log_10_(export) = *m*×log_10_(NPP)+*b*, with *m* = 0.76 ± 0.1 and *b* = −0.08 ± 0.3. Since this slope is less than 1 it implies that the export ratio (panel b) is inversely proportional to net primary production. d) Anomaly relative to the mean (in percent change) of one-year running means of net primary production estimated by satellite remote sensing (green, Kahru et al. 2015), sinking carbon export estimated from the model of Kelly et al. (2018), which uses a regression of sediment trap-based export on NPP (red), and sinking carbon export from the model of Messié et al. (2025), which is forced using nutrient supply estimates. All regressions are Type II geometric mean regressions.

While this offshore advection partially resolves the spatial discrepancy between new and export production, it leaves open the question of how strongly the gravitational pump will respond to temporal changes in net primary production (Fig. 4d). Two distinct approaches have been developed for using satellite-based measurements to estimate export flux patterns in the CCE. The first approach is based on regressions of sediment-trap-measured gravitational carbon export on net primary production (Stukel et al. 2015b, Kelly et al. 2018, Kahru et al. 2020).

Because they are based on the inverse correlation between export efficiency and net primary production in Fig. 4b, these estimates predict that export will exhibit a proportionally weaker temporal variability than net primary production (Fig. 4d). The other approach for regional satellite-based carbon export estimates combines estimates of regional nitrate supply (i.e., potential new production) with satellite-observed surface currents (Messié et al. 2025). Because this approach is fundamentally driven by nitrate supply, it shows a similar response to new production (Fig. 4a) and predicts proportionally stronger temporal variability for export production, relative to net primary production (Fig. 4d). The discrepancy between these two approaches is exemplified by focusing on one of the strongest perturbations to the ecosystem in the observational record, the 2014-2015 “Blob” marine heatwave, which was immediately followed by the 2015-2016 El Niño. Export estimates using the sediment trap regressions suggest that the gravitational pump declined by only ∼10% (Fig. 4d, red line). However, export estimates forced with upwelled nutrient supply suggest that the BCP decreased by nearly 50% during the peak of this event (Fig. 4d, blue line). This contradiction might arise because the “space-for-time” approach breaks down in advective systems with long lag times, suggesting that the gravitational pump responds similarly to new production (Fig. 4d, blue line). However, an alternate hypothesis exists. While the sum of all BCP pathways must be equal to new production, it is not necessarily true that the gravitational pump should respond identically to new production. The alternate hypothesis thus suggests pathway flexibility; during periods of low productivity, the gravitational pump dominates the BCP while during high productivity periods subduction and/or active transport become more important. Understanding how the BCP and carbon sequestration within EBUSs will respond to predicted changes in climate thus necessitates greater understanding of these processes.

### The ecology of sinking particles: Production

Microscopic examination of sinking particles exiting the euphotic zone show a dominant role for mesozooplankton fecal pellets, especially in the coastal domain (Fig. 5, Stukel et al. 2013, Morrow et al. 2018). These results are supported by pigment measurements, which show that concentrations of phaeopigments – a product of chlorophyll degradation within zooplankton guts – are much higher than chlorophyll *a* concentrations (Landry et al. 1994, Morrow et al. 2018).

**Fig. 5.**
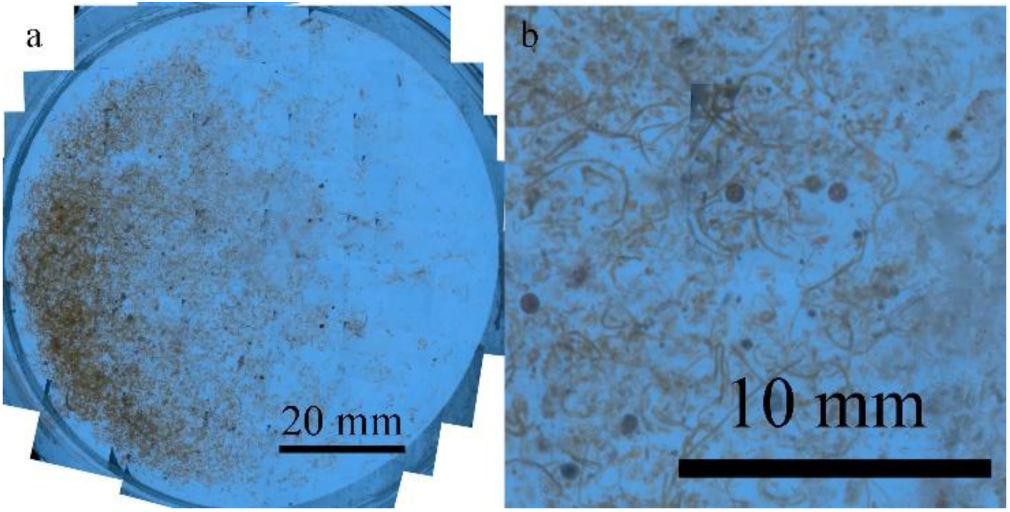
Sinking particles collected in a sediment trap deployed at 50-m depth beneath a coastal region of the CCE. Figure modified from Fender et al. (2019).

**Fig. 6.**
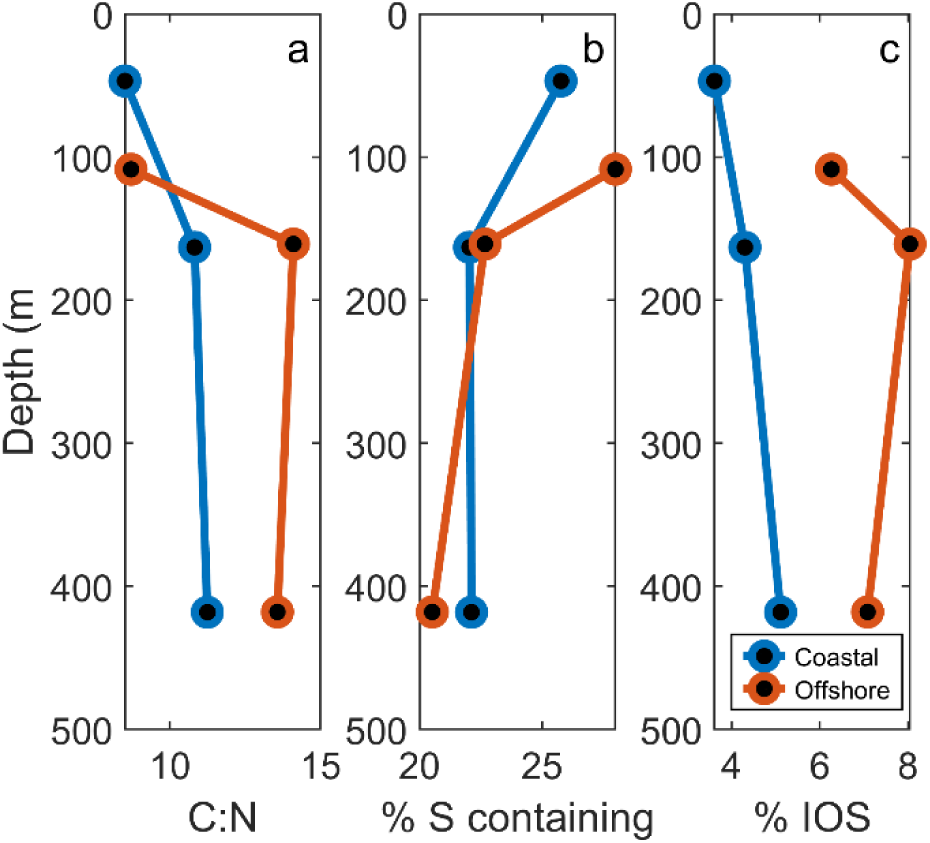
Molecular characteristics of sinking organic matter at a coastal upwelling (inshore) and oligotrophic offshore site. a) bulk organic carbon:nitrogen ratio (mol:mol). b) relative abundance (%) of S-containing molecules in the water-extractable fraction of the sinking particles as measured by Fourier-transform ion-cyclotron-resonance mass spectrometry (FT-ICR-MS). c) relative abundance (%) of “Island of Stability” (IOS) molecules in the water-extractable fraction measured by FT-ICR-MS. IOS molecules are a stable class of molecules consistently found in deepwater dissolved organic matter (Lechtenfeld et al. 2014). Data from Forrer et al. (in review),

Copepods and euphausiids (krill) were the dominant taxa responsible for creating these pellets, with copepods contributing a relatively constant carbon flux, while euphausiid contributions were highly variable. Other taxa contribute as well, with salps and anchovies occasionally producing millimeter-scale and rapidly sinking fecal pellets that likely contribute to deep ocean carbon storage (Stukel et al. 2013, Smith et al. 2014, Fender et al. 2019, Bourne et al. 2021), although their contribution is sporadic. Further, the impact of a recent decade-long increase in abundance of the colonial pelagic tunicate *Pyrosoma atlanticum* is potentially important but has yet to be assessed (Miller et al. 2019). Across all of the pellets collected in the traps, the median (carbon-weighted) size was estimated at an equivalent spherical diameter of 265 µm, which equates to an estimated median sinking speed of 141 m d^-1^ (Stukel et al. 2019b). While fecal pellets clearly dominate export flux in the coastal upwelling region, their proportional contribution to carbon flux declines in the offshore region and appears to decrease with depth (Morrow et al. 2018, Bourne et al. 2021). In these offshore and deeper regions, flux was typically dominated by the gravitational sinking of amorphous marine snow aggregates of unknown origin. Despite the strong evidence that zooplankton fecal pellets are the dominant type of sinking particle exiting the euphotic zone in the CCE, no correlation has been found between gravitational pump efficiency and zooplankton abundance or biomass.

While pigments suggest that intact phytoplankton are a relatively minor contributor to sinking flux in the region (Morrow et al. 2018), studies have nevertheless linked specific phytoplankton taxa to periods of high export flux (Preston et al. 2019, Valencia et al. 2021).

Specifically, the diatom *Thalassiosira* was conspicuously dominant during a high pulse of export measured on the bathypelagic seafloor (Preston et al. 2019) and also the only phytoplankton taxon that was consistently enriched in shallow sediment traps relative to its presence in the euphotic zone community (Valencia et al. 2021, Valencia et al. 2022). Importantly, while the large, dense siliceous frustules of this taxon suggest that it is a likely contributor to export, it has also been shown to be digestion resistant (Fowler and Fisher 1983), suggesting that its overrepresentation in sediment trap DNA sequences may be a result of a preservational bias within zooplankton fecal pellets (Valencia et al. 2021). Carbon export roles by diatoms have also been shown to depend on the physiological status of cells - Fe-limitation induces increased silicification and commensurately increased density and sinking speeds for sinking particles – although the dense diatoms are likely incorporated into zooplankton fecal pellets prior to sinking (Brzezinski et al. 2015, Stukel et al. 2017, Stukel and Barbeau 2020).

Other phytoplankton taxa are at times important to export flux, as identified from DNA sequencing studies, including some dinoflagellates, chlorophytes, and *Synechococcus* (Preston et al. 2019, Valencia et al. 2022). Importantly, *Prochlorococcus*, the most abundant phototroph in the world ocean and in offshore regions of the CCE (Chisholm et al. 1988, Flombaum et al. 2013, Taylor and Landry 2018), is rare in sediment trap samples and almost certainly underrepresented relative to its contribution to the overlying water column community (Valencia et al. 2021). Rhizarians – and especially polycystine radiolarians and phaeodarians – have also been found to be important contributors to export flux in the CCE (Biard et al. 2018, Gutierrez-Rodriguez et al. 2019, Preston et al. 2019, Valencia et al. 2022). Like diatoms, these taxa are silicified and hence substantially denser than many other plankton, although their ecological roles as heterotrophs and/or mixotrophs are distinctly different than those of diatoms (Gowing 1986, Caron et al. 1995, Suzuki and Not 2015, Boltovskoy et al. 2017). Substantial interannual variability in the abundances of Rhizaria suggests varying roles in the BCP (Stukel et al. 2018a, Biard and Ohman 2020).

Beyond phytoplankton, evidence is emerging that the microbial communities associated with sinking particles are distinct relative to those in the water column. Valencia et al. (2022) sampled sinking particle and upper water column communities across a productivity gradient spanning a seven-fold range in vertically integrated primary production and a sixty-fold range in surface chlorophyll. Across this large range of environmental conditions, a dramatic shift was seen in phytoplankton communities from diatom-dominated near the coast to *Prochlorococcus*-dominated in the offshore nutrient-depleted region. Nevertheless, 16S rRNA sequencing of prokaryotic communities suggested that sinking particle communities from offshore or coastal communities were more closely related to each other than either was to the local water column community. Results from 18S rRNA sequencing of eukaryotes were similar, although slightly more nuanced, suggesting that eukaryotic communities associated with sinking particles resemble the overlying community more than prokaryotic communities. Similarly, Gutierrez-Rodriguez et al. (2019) found that eukaryotic communities on sinking particles were distinct from euphotic zone communities, while Preston et al. (2019) found that specific eukaryotic and prokaryotic taxa (including Alteromonodales and Flavobacteriales) were present in all deep-ocean sediment trap samples collected over a nine-month period. These results suggest that some taxa may play key roles in the gravitational pump, although it is not necessarily clear which taxa are responsible for contributing to sinking particles, colonizing those sinking particles, and/or simply over-represented on sinking particles because of characteristics that promote preservation of their genetic material (Valencia et al. 2021).

Research in the CCE has also identified the importance of enhanced carbon export in the vicinity of mesoscale features including fronts (Brzezinski et al. 2015, Stukel et al. 2017) and coastal filaments (Kelly 2020, Kranz et al. 2020). These regions are sites where intense vertical mixing can introduce nutrients to the surface ocean, while lateral mixing between water parcels with different limiting nutrients can further stimulate algal growth (Franks 1992, Li et al. 2012). Biomasses of phytoplankton and zooplankton are often enhanced at fronts and filaments as a result of both in situ growth and passive transport (Nagai et al. 2015, Powell and Ohman 2015, de Verneil et al. 2019, Gangrade and Franks 2023). They are thus sites in which the BCP is enhanced both through increased production of sinking particles and increased downward flux of non-sinking organic matter (both dissolved and particulate) due to subduction and vertical mixing (Stukel et al. 2017). The inherent heterogeneity of these systems may also drive nonlinear dynamics as consumers exploit rare resource patches. Large-scale-climate driven changes to the frequency of frontal systems in the CCE (Kahru et al. 2012, Kahru et al. 2018) further suggest that the BCP may respond to climate-driven shifts in mesoscale dynamics.

### The ecology of sinking particles: particle transformation and breakdown

The amount of carbon sequestered by the BCP depends not only on how much organic carbon is exported out of the surface ocean, but also on the depth to which that carbon penetrates before it is remineralized back to CO_2_. Indeed, Kwon et al. (2009) estimated that a 24-m increase in the mean depth of remineralization globally could lead to a drop in atmospheric CO_2_ of 10 – 27 ppm. Estimates of carbon flux attenuation (i.e., the decrease in carbon export with depth in the ocean) are highly variable across the world ocean (Marsay et al. 2015), and likely depend on a suite of ecological and environmental parameters including temperature, oxygen, and community composition (Cavan et al. 2017, Laufkötter et al. 2017). Carbon flux attenuation is often quantified using either Martin’s *b*, a key parameter from the Martin curve (where *b* is the negative exponent of a power law relationship between carbon flux and depth, and a higher *b* denoting increasingly rapid flux attenuation; Martin et al. 1987) or transfer efficiency (T_100_ = carbon flux at a depth 100 m deeper than the euphotic zone depth divided by carbon flux at the euphotic zone depth; Buesseler and Boyd 2009). Within the CCE, flux attenuation with depth is typically moderate. Stukel et al. (2023) estimated an average regional *b* of 0.72 (for comparison typical *b* values globally range from 0.4 - 1.6, Marsay et al. 2015) and T_100_ values averaged a modest 0.54 (for comparison the global compilation of Buesseler et al. 2020 found a mean T_100_ = 0.68, st.dev. = 0.33). Together, these results indicate that although this region is highly productive, it is only moderately efficient at exporting carbon to depth.

These net carbon flux attenuation metrics are the result of complex ecological interactions that transform, modify, re-package, disaggregate, solubilize, and remineralize sinking particles. Sediment trap nucleic acid analyses clearly identify specific taxa that are associated with sinking particles. Some, such as the prokaryotes Alteromonodales and Flavobacteriales are known to colonize particles and hence likely play roles in breaking down these sinking particles (Preston et al. 2019). Others, including the heterotrophic nanoflagellate *Caecitellus parvus*, increase in abundance in un-preserved sediment trap samples suggesting that they are actively feeding either on the particles themselves or on the associated microbial communities. These results depict a system in which rapid colonization of sinking particles stimulates community succession as the particles sink through the mesopelagic, potentially reducing export efficiency and reshaping the molecular stoichiometry of sinking particles.

Changing particle dynamics with depth are also evidenced in broader characteristics of the sinking particles. For instance, the large phaeodarian *Aulosphaeridae* (giant siliceous protists that are ∼2-mm in diameter) were relatively negligible contributors to sinking flux leaving the base of the euphotic zone, but substantial contributors at a depth of 150 m (Biard et al. 2018). These taxa are flux feeders believed to consume sinking particles and typically live at 100 – 150 m depth in the CCE (Stukel et al. 2018a, Biard and Ohman 2020). Their presence in the sediment traps thus implies that they consumed sinking particles, transformed the carbon into their own biomass, and then after their deaths contributed further to deeper flux into the mesopelagic. Similarly, results from beneath a coastal filament suggested a shift from predominantly fecal pellets of anchovies and zooplankton at shallow depths to aggregates (possibly appendicularian houses) in the mesopelagic (Bourne et al. 2021). This highlights a diverse range of particle transfer efficiencies to depth: while some sinking particles undergo multiple transformations as they travel to depth, others – often rapidly sinking and derived from near-surface-derived origins (e.g., salp fecal pellets and carcasses) – have been identified on the seafloor or in near-bottom sediment traps virtually intact, indicating minimal diagenesis (Smith et al. 2014).

Changes are also reflected in the molecular characteristics of sinking particles. The C:N ratio of sinking particles exiting the base of the euphotic zone in the CCE is typically slightly higher than the canonical “Redfield ratio” of 6.6 mol:mol (Redfield 1934), with a median of 7.5 mol:mol (although with substantial variability, st.dev. = 1.9). The C:N ratio generally increases with depth (median at ∼150-m depth = 8.0, median at ∼450-m depth = 11.6 mol:mol), reflecting preferential microbial utilization of nitrogen-rich compounds such as amino acids. Forrer et al. (in review) determined that the proportion of molecules with elemental ratios similar to those that have been found to be enriched in the deep ocean DOM increased with depth. Sulfur-depleted and recalcitrant molecules were also more common beneath the oligotrophic areas of the CCE than beneath the coastal upwelling domain. Taken together, these results suggest that labile organic molecules are preferentially utilized on sinking particles and hence the organic matter supplied to the deep ocean is likely less bioavailable than that at shallower depths. Forrer et al. (in review) also suggests that particle degradation and transformation begins within the euphotic zone, the biogeochemical dynamics of which create both a vertical and horizontal gradient in the quality of sinking particulate organic matter (Fig. 7).

**Fig. 7.**
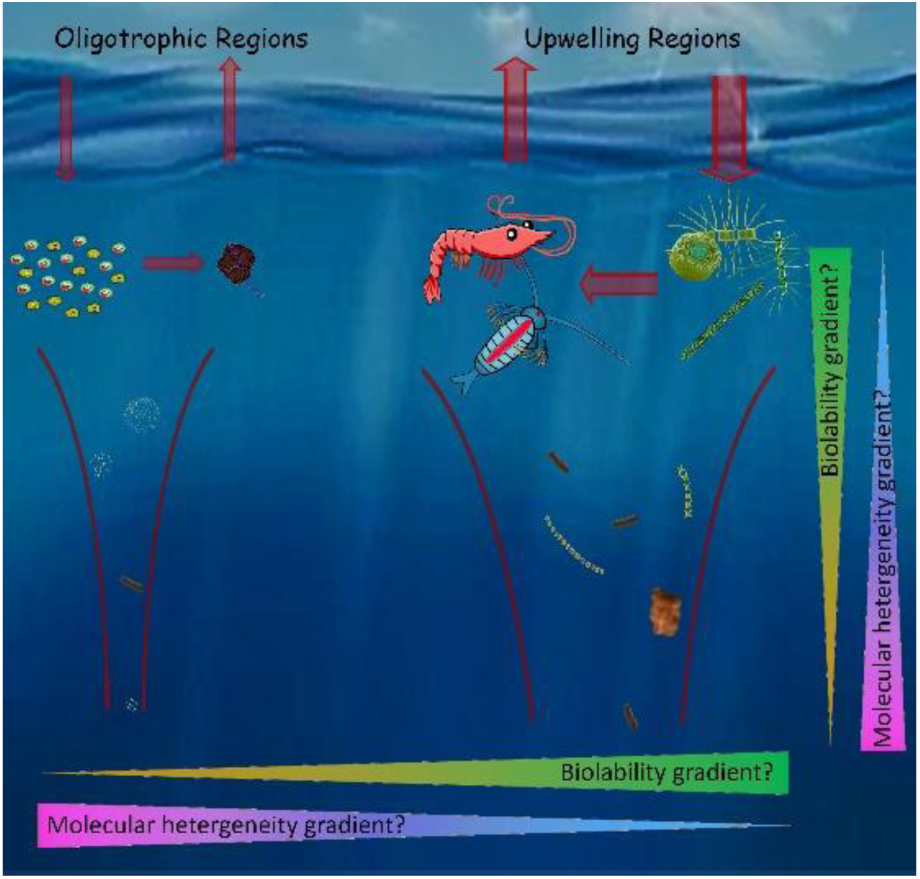
A conceptual model for molecular characteristics of the biological carbon pump. Biolability (the relative ease with which molecules can be used by marine microbes) is hypothesized to decrease with increasing depth and in regions with less nutrients. In contrast, molecular heterogeneity (the diversity of different molecules) increases with increasing depth and in low nutrient regions.

In addition to particle-attached microbes, a wealth of deep-ocean metazoans rely on sinking carbon, either directly or indirectly. Stukel et al. (2019b) estimated that suspension-feeding zooplankton could consume the majority of slowly sinking (<10 m d^-1^) particles before they could sink through the upper 200 m of the water column, although these suspension-feeders had minimal impact on the rapidly sinking (>100 m d^-1^) particles that are assumed to dominate flux. In contrast, two “flux feeding” zooplankton (i.e., zooplankton that feed by passively intercepting sinking particles), the phaeodarian *Aulosphaeridae* and the pteropod *Limacina helicina*, might each be responsible for ∼10% of measured carbon flux attenuation in the upper twilight zone (Stukel et al. 2019b). The total demand of mesopelagic zooplankton and micronekton in the CCE has been estimated to range from 75 – 464 mg C m^-2^ d^-1^ (Kelly et al. 2019). Numerical and conceptual models often treat these deep ocean communities simply as consumers of sinking particles. However, observational, isotopic, and modeling studies in the CCE all depict a much more complex mesopelagic food web with distinct predator-prey relationships and multiple trophic levels supported by sinking particles (Choy et al. 2017, Kelly et al. 2019, Stukel et al. 2019a, Hetherington et al. 2024).

The impact of metazoan communities on sinking particles likely varies with depth and between different EBUSs due to differences between the oxygen minimum zones that underly these regions (Karstensen et al. 2008, Seibel 2011). The Humboldt Current overlies the strongest oxygen minimum zone, with oxygen becoming limiting slightly beneath the euphotic zone and typically become truly hypoxic at depth ranges from ∼150 – 700 m (Fig. 8). The CCE also overlies a strong oxygen minimum zone, in part because of its proximity to other low oxygen waters (the Eastern Tropical Pacific and the North Pacific Subtropical Gyre). However, because of colder surface temperatures and lower productivity (relative to the Humboldt Current) oxygen concentrations typically do not become limiting until ∼300 m depth and only become truly hypoxic at depths >500 m. Oxygen minimum zones beneath the Canary and Benguela EBUSs are weaker, although oxygen levels beneath the Canary EBUS drop to levels that can be potentially limiting for metazoans much more rapidly than they do in the CCE (Fig. 8). These low oxygen conditions can limit respiration by both microbial and metazoan communities thus increasing carbon transfer into the deep ocean (Seibel 2011, Ulloa et al. 2012, Cavan et al. 2017, Weber and Bianchi 2020, Busecke et al. 2022). As a result of climate change, oxygen minimum zones in all of these regions are expanding (Bograd et al. 2008, Keeling et al. 2010, Gilly et al. 2013, Busecke et al. 2022), while synergistic effects between oxygen and temperature suggest that biotic responses will be substantial (Deutsch et al. 2015).

**Fig. 8.**
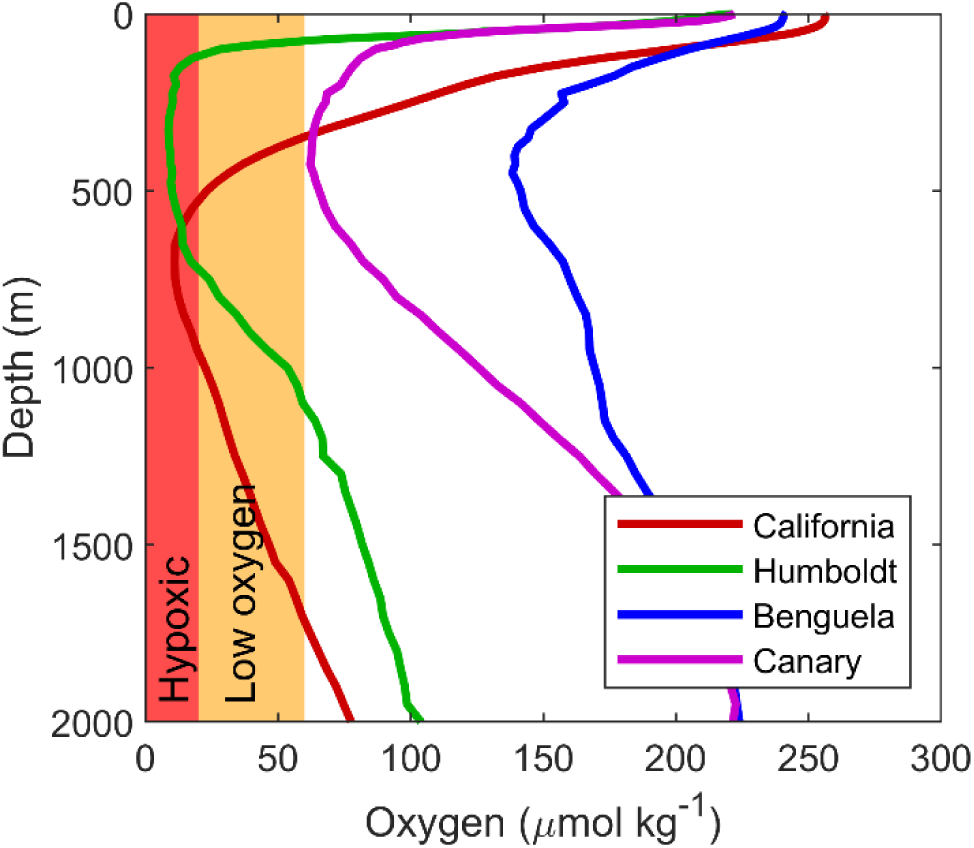
Typical oxygen profiles in the major EBUSs. Data from Garcia et al. (2024)

### Active transport by vertical migrants

Many zooplankton, small fish, and squid vertically migrate in synchrony each day; rising at twilight and descending to depths of 200 – 600 m at dawn (Cushing 1951, Steinberg et al. 2000, 2013, Archibald et al. 2019). This strategy minimizes risk of predation by visual predators, while also potentially providing bioenergetic, reproductive, or transport advantages (Ohman 1990, Hays 2003, Bandara et al. 2021). By feeding in the euphotic zone before descending to depth, these vertical migrants contribute to downward carbon flux through three distinct mechanisms: 1) respiration and excretion at depth, 2) fecal pellet production at depth, and 3) mortality at depth (Longhurst and Harrison 1989, Steinberg et al. 2000). Respiration and excretion at depth of carbon consumed in the euphotic zone are often assumed to be most important. While they are not frequently measured directly, they can be estimated from day-night biomass differences combined with allometric- and temperature-based estimates of respiration rates in the mesopelagic zone (Ikeda et al. 2007). Fecal pellet production at depth (i.e., defecation in the deep ocean of organic matter consumed in the surface ocean) may be important in cold water regions but is not likely to be important in the CCE where relatively warm surface temperatures lead to gut turnover times on the order of 1 h for zooplankton (Dam and Peterson 1988). In contrast, mortality rates for zooplankton and nekton in the deep ocean are highly uncertain. Indeed, few studies have been able to either follow cohorts of these taxa for sufficiently long periods to accurately estimate mortality or observe statistically significant numbers of predator-prey interactions even in the surface ocean (Ohman and Hirche 2001, Ohman and Hsieh 2008, Thiebot et al. 2016, Choy et al. 2017). Predation risk is likely highly variable for zooplankton and nekton and varies in response to the predator field, the light environment, and prey behavior and morphology (Aksnes and Giske 1993, van Someren Gréve et al. 2017).

In the CCE paired day-night net tows, day-night differences in zooplankton acoustic backscatter measured by autonomous gliders, and multi-frequency shipboard acoustic backscatter of midwater fishes have all been used to estimate the biomass of vertical migrators (Stukel et al. 2013, Davison et al. 2015, Powell and Ohman 2015). Copepod vertical migration behavior has been shown to be size dependent, with 2 – 8 mm copepods migrating the deepest (average amplitude of daily migrations of ∼100 m), presumably because smaller copepods have lower visual predation risk and hence can remain near the surface, while larger copepods are visible even under dim illumination and hence remain at depth (Ohman and Romagnan 2016). Estimates of euphausiid (krill) vertical migration are more variable as a result of the patchiness of these organisms, although they are often responsible for the greatest active transport and typically migrate from the euphotic zone to depths of 200 – 500 m (Stukel et al. 2013).

Myctophids (∼5-cm midwater fish also known as lanternfish) are another important vertical migrator with typical daytime residence depths of ∼450 m (Davison et al. 2013, Netburn and Koslow 2015). Respiratory fluxes associated with zooplankton vertical migration have been estimated to transport 2.1 (95% C.I. = 1.1 – 3.3) mmol C m^-2^ d^-1^ out of the upper ocean (Stukel et al. 2013, Stukel et al. 2023), with myctophids and other midwater fish adding an additional ∼0.8 mmol C m^-2^ d^-1^. While the stoichiometry of active transport has not been estimated in the CCE region, measurements from the contiguous North Pacific Subtropical gyre suggest that vertical migrant export is associated with lower C:N and C:P ratios than that of sinking particles (Hannides et al. 2009).

The carbon flux associated with mortality of these organisms at depth is highly uncertain, however, with CCE regional estimates ranging from mortality at depth being a small fraction of total export to approximately equal to respiratory losses (Kelly et al. 2019, Stukel et al. 2022). Video from remotely operated vehicles have identified multiple midwater predators that feed on zooplankton and micronekton as they migrate, including ctenophores, siphonophores, and squid (Robison et al. 2020). Given correlations between the populations of such predators and their vertically migrating prey (Koslow et al. 2014), it is reasonable to surmise that mortality risk is positively correlated with vertical migrator biomass and Stukel et al. (2022) estimated that daily mortality rates in the deep ocean could range from 0.3% d^-1^ in oligotrophic areas to potentially as high as 6% d^-1^ in high biomass regions.

### Subduction and vertical mixing

Carbon export via subduction and vertical mixing are notoriously difficult to measure with observations, because it is driven by physical circulation that varies across many spatial and temporal scales. Boyd et al. (2019) summarized the portions of the BCP that transport particles to depth and subdivided the physically driven portion into the eddy-subduction pump (driven by meso- and submesoscale eddies and fronts), the large-scale subduction pump (which is determined by Ekman pumping and horizontal circulation across ocean basins), and the seasonal mixed layer pump (driven by deepening of the mixed layer in wintertime at mid- to high latitudes). The first two are likely most important in the CCE, where the seasonal mixed layer remains shallow throughout the year. Bograd and Mantyla (2005) searched for deepwater oxygen and nutrient signatures of subduction and found these to be common and persistent in the CalCOFI observational record, often at depths of ∼200 m. These signatures are likely evidence of rapid export via the eddy-subduction pump, which was further found to be particularly important near mesoscale eddies, where subduction can be as important as sinking particle flux (Stukel et al. 2017). The importance of the eddy-subduction pump can be overestimated by such methods, however, because downward flux along mesoscale eddies and fronts is often balanced by upward flux in other regions of the mesoscale feature leading to substantially reduced net carbon transport (Resplandy et al. 2019). Combining the effects of the eddy-subduction and large-scale subduction pumps, downward particulate organic carbon transport has been estimated to be approximately equal to gravitational flux at the 100-m depth horizon, but to decline substantially with depth, such that subduction sequesters less than ¼ of the amount of carbon sequestered by gravitational flux (Stukel et al. 2018b, Stukel et al. 2023). The above estimates do not, however, include the additional transport mediated by dissolved organic matter. Lateral advection of dissolved organic matter from the coast to offshore has been shown to play an important role in the explaining the spatial decoupling of new and export production (Stephens et al. 2018), although the persistence time scales for this organic matter during mesoscale or large-scale subduction events is not known. Importantly, the C:N and C:P ratios for dissolved organic molecules are typically higher than those for sinking particles suggesting that if dissolved organics are a primary component of the subduction pumps then these physical pathways may export more carbon per unit nitrogen than the gravitational pump (Hopkinson Jr and Vallino 2005, Zakem and Levine 2019, Liang et al. 2023).

### Climate change and carbon sequestration in coastal upwelling ecosystems

Predicted climate change impacts on EBUSs are complex (Bograd et al. 2023). Upwelling-favorable winds are driven by large-scale atmospheric pressure gradients between the subtropical ocean basins and adjacent land. These winds intensify when the continents warm in summer and hence Bakun (1990) hypothesized that anthropogenic warming would lead to stronger coastal upwelling. Results from climate simulations have further highlighted that poleward migration of the subtropical ocean high-pressure zones will also modify the location and timing of upwelling (Rykaczewski et al. 2015). The impact of these upwelling favorable winds, however, may be offset by increased stratification, especially in the offshore extent of EBUSs (Jacox and Edwards 2011). Indeed, results from 73-years of optical clarity measurements (used as a proxy for phytoplankton biomass) in the CCE suggest that coastal regions have become progressively more productive while the offshore domain has become more oligotrophic (Kahru et al. 2023). The impacts of climate change on phytoplankton production is further complicated because remote changes in the nutrient concentrations of source waters for upwelling may alter nutrient supply irrespective of changes to upwelling (Rykaczewski and Dunne 2010, Bograd et al. 2015, Jacox et al. 2024). Recent evidence from global climate simulations further complicate predictions by suggesting that altered geostrophic flows (i.e., ocean circulation driven by large-scale differences in sea surface height and ocean density structure) might offset stronger upwelling-favorable winds (Jing et al. 2023). These impacts may be most pronounced in the two southern hemisphere EBUSs; the Humboldt and Benguela systems may actually warm faster than surrounding waters (Wang et al. 2023). As this review is focused on the BCP, rather than ocean circulation, we simply note here that EBUS physical and nutrient-supply responses to climate change are complex and will drive modifications to the BCP; understanding whether these changes will increase or decrease net CO_2_ uptake by EBUSs requires a better understanding of how export efficiency and stoichiometry vary temporally in response to changing drivers.

While studies highlighted in previous sections give us substantial insight into the physical and biogeochemical processes that drive BCP patterns in the CCE, it is less clear how these intersecting processes will control interannual variability in export flux and how the regional BCP will respond to a changing climate (e.g., Fig. 4d). Results from deep-sea sediment traps show substantial variability in sinking organic matter reaching the seafloor (Fig. 9, Sekula-Wood et al. 2012, Smith et al. 2013). These results show sensitivity of organic carbon flux to climatic drivers and to shifts in the zooplankton community (Smith et al. 2006, Smith et al. 2013, Smith et al. 2014). Within the Santa Barbara Basin (∼500-m depth) deep-sea sediment traps have shown a correlation between sinking organic carbon flux and sinking mineral flux (carbonate, silica, and lithogenic material), suggesting that the ballasting effect associated with increased mineral content is important in determining flux into the deep ocean (Thunell et al. 2007).

**Fig. 9.**
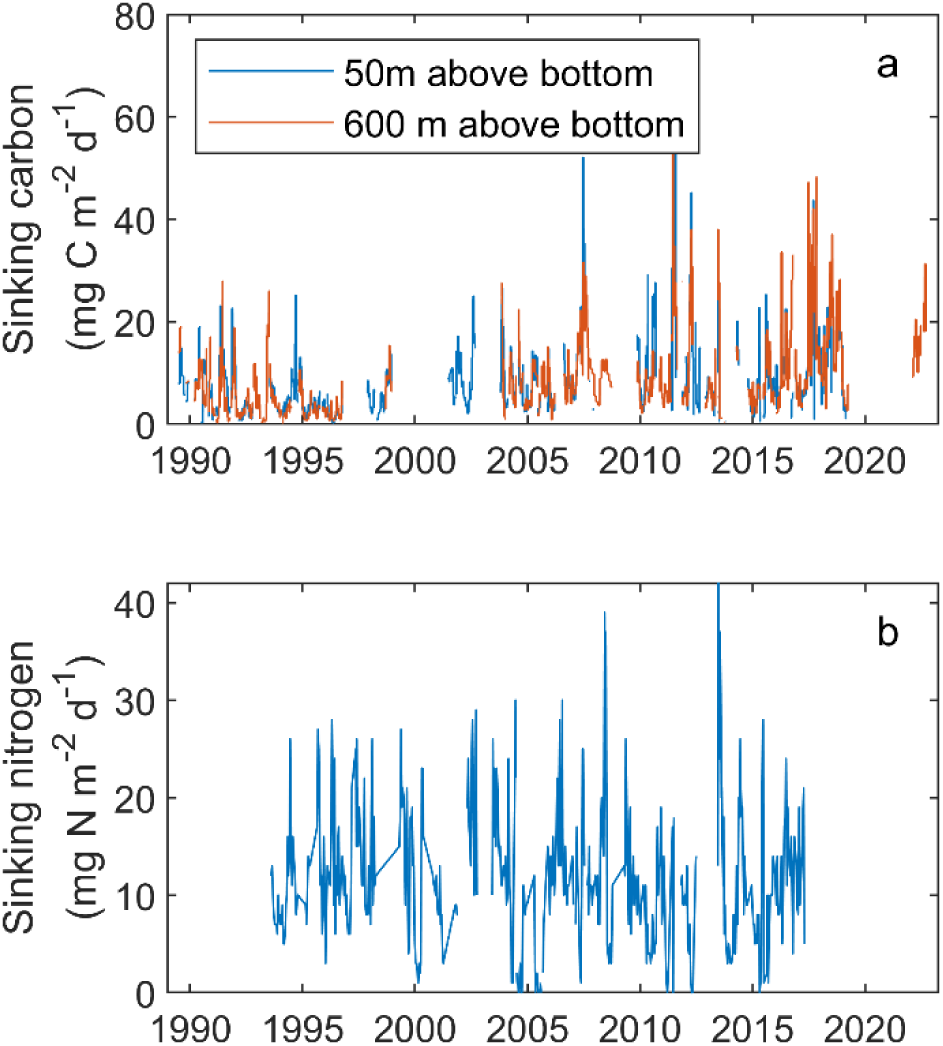
Sinking organic matter flux into deep-sea moored sediment traps. a) Sinking organic carbon flux above the abyssal plain at ∼4000 m depth in an offshore region of the California Current Ecosystem. Two traps are typically deployed at this station with one 50 m above the bottom (∼3950-m depth) and the other 600 m above the bottom (∼3400-m depth) (Smith et al. 2018, Smith et al. 2020). b) Sinking nitrogen flux into a sediment trap in the Santa Barbara Basin, a much more productive and shallower (sediment trap depth is ∼500 m) region of the California Current Ecosystem (Thunell et al. 2007, Sekula-Wood et al. 2012).

Nevertheless, sinking silica fluxes at this site are not strongly correlated with silicon dynamics of surface communities (Shipe et al. 2002) and a surprising increase in sinking organic carbon flux was observed during the 1997-1998 El Niño (Shipe et al. 2002). Comparison to surface phytoplankton communities have further elucidated distinct differences, with, e.g., warm-water surface diatoms seldom found in deep-sea sediment trap samples (Venrick et al. 2003). Results at this site also show high organic C:P ratios for sinking particles, suggesting rapid remineralization of phosphorus (Sekula-Wood et al. 2012).

At the Station M study site, which is in substantially deeper water (∼4000-m depth) in the deep-ocean abyssal plain, a 30-year time series has demonstrated long-term variability in sinking carbon flux to the seafloor linked to climate drivers including El Niño-Southern Oscillation and the Northern Oscillation Index (Smith et al. 2006). These studies demonstrated that under typical conditions, sinking carbon flux is insufficient to meet the carbon demand of the benthic community, although this deficit may be explained by episodic pulses of particle export (Smith et al. 2013). Further, a long-term increase in the proportion of total flux to the benthos mediated by episodic pulses of sinking materials at ∼4000-m has been observed (Smith et al. 2018). These pulses have been linked to coastal diatom taxa, suggesting that variability in flux at this deep-sea site is related to offshore currents linking coastal production to the deep-ocean abyssal plain (Preston et al. 2019, Michaud et al. 2022), as well as to stochastic salp blooms (Smith et al. 2014). Relating deep-sea carbon flux variability to surface ocean biology has been challenging, likely as a result of the aforementioned lateral transport processes combined with temporal lags that can vary as a result of different particle sinking speeds and processing by midwater biota (Kahru et al. 2020, Ruhl et al. 2020, Messié et al. 2023, Messié et al. 2025).

While these results demonstrate links between climate drivers and deep-sea particle flux, they also demonstrate that these deep-sea sediment traps are not sufficient for quantifying regional variability in particle flux at the surface, both because they reflect single-location measurements, and because multiple processes can alter subsurface flux attenuation patterns. One particularly important subsurface process is the growth of the oxygen minimum zone (Bograd et al. 2008). Increased hypoxic volume will directly retard microbial remineralization on sinking particles (Van Mooy et al. 2002, Ulloa et al. 2012, Engel et al. 2017). The depth of the deep scattering layer (a mesopelagic layer with high zooplankton and micronekton biomass) may also shoal and thicken as a result of climate change and ocean deoxygenation (Netburn and Koslow 2018), leading to increased consumption of particles in the shallow twilight zone.

Indeed, an increase in the oxygen minimum zone is already causing a decline in CCE mesopelagic fish communities (Koslow et al. 2011).

### A plug-flow-reactor conceptual model for biological carbon pump responses

Given the absence of reliable time-series of carbon export from the upper ocean via sinking particles, active transport, and subduction, it is useful to consider a conceptual model for how these processes might vary. We suggest a plug-flow-reactor conceptual model as an organizing framework. With this model, an initial pulse of upwelling will dilute the surface ocean with cold, CO_2_- and nutrient-rich, POC- and DOC-depleted deep water while also beginning to push the affected water parcel offshore via Ekman transport. Plankton communities will respond to this upwelling pulse through a predictable successional pattern with rapid (i.e., ∼one week) phytoplankton bloom formation and a lagged (two to four-week) increase in zooplankton abundances (Stukel et al. this issue). Since sinking particle flux in EBUSs is typically comprised mostly of zooplankton fecal pellets (and zooplankton grazing is ∼proportional to zooplankton biomass × phytoplankton biomass) it is reasonable to assume that sinking carbon flux will peak earlier in the successional pattern than active transport (which is ∼proportional to the biomass of vertically migrating zooplankton). Eventually, the entire water parcel will get subducted from the surface ocean, at which point it will lose contact with the atmosphere (no further gas exchange) and organic matter remaining in the water parcel can be considered sequestered.

Under this conceptual model, the relative balance between sinking, active transport, and subduction pathways are largely determined by how long the water parcel remains at the surface before it is subducted; very short durations would indicate that subduction dominates, moderate durations would see sinking dominate, and longer durations would exhibit growing importance of active transport (Fig. 10). Since subduction must balance upwelling (although typically laterally displaced), upwelling-favorable conditions, e.g., La Niñas or typical spring conditions, will feature short surface durations for water parcels with greater importance for subduction.

**Fig. 10.**
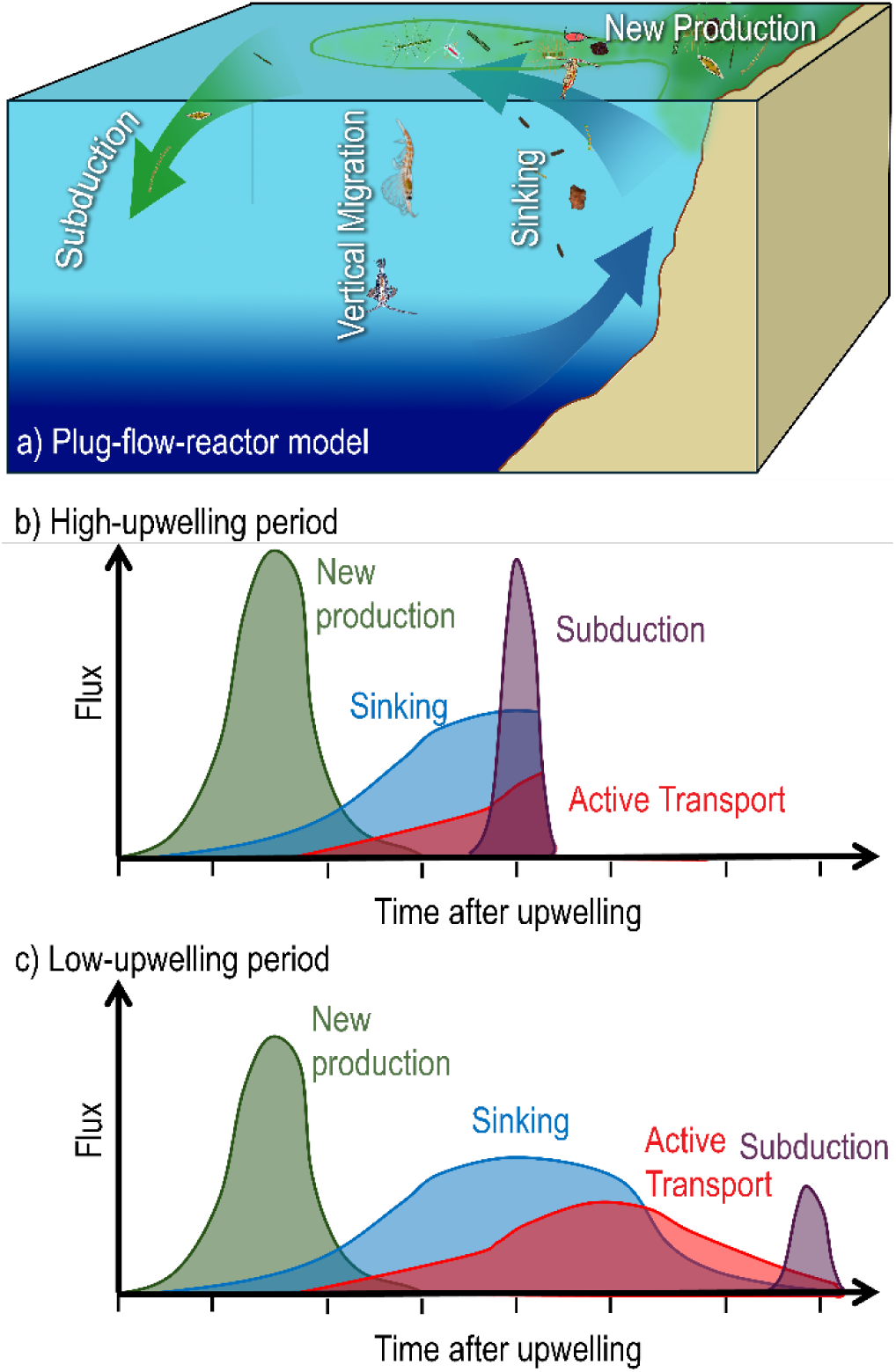
Plug-flow-reactor conceptual model. a) Upwelling introduces nutrients into the surface ocean, stimulating new production and a phytoplankton bloom, while also causing Ekman transport to push the water parcel away from the coast. The ultimate fate of the water parcel is subduction, but along the way the plankton community will undergo a successional cascade in which the phytoplankton bloom exhausts nutrients and a community of zooplankton grazers crops down the phytoplankton community. New production will typically peak early in this cascade with peaks in sinking particle flux and active transport by vertical migrants occurring later. b) When upwelling rates are high, the water parcel may not be at the surface long enough for this successional cascade to play out and there will still be substantial organic matter still remaining in the water parcel when it is subducted. c) When upwelling rates are slow, the water parcel may remain in the surface for a longer period of time, allowing sinking particle flux and vertical migration to remove most of the organic matter before it is subducted.

Conversely, upwelling-depressed periods, e.g., El Niños, will have longer surface durations and see greater importance for sinking and active transport. Ultimately, since the amount of upwelled nitrate must balance the amount of exported nitrogen within organic matter (Eppley and Peterson 1979), *net* biologically mediated uptake of CO_2_ through this sequence will be determined by the relative C:N stoichiometry of upwelled inorganics (total CO_2_:NO_3_^-^) and exported material. If the C:N ratio of exported material remains the same over time, then current rates of nutrient supply will support enough export production to exactly balance the amount of CO_2_ upwelled with the nutrients (since this CO_2_ was created during the same organic matter decomposition that released nutrients into the deep ocean). However, if the C:N ratio of exported material increases, the BCP will sequester more CO_2_ in the future, while if the C:N ratio decreases it will sequester less. Importantly C:N ratios are likely the highest for subducted organic matter (due to the contribution of high C:N dissolved organic matter), followed by sinking particles (typical C:N ratios are 6 – 10 based on shallow CCE LTER sediment traps) and active transport (C:N = 4.25, Hannides et al. 2009). This plug-flow-reactor conceptual model thus suggests that upwelling favorable conditions will lead to net CO_2_ uptake within EBUS’s, while upwelling-depressed conditions will lead to net CO_2_ efflux.

The above analysis is a crude approximation for a complex system. It does not account for some key processes, such as increased denitrification within oxygen minimum zones which may alter the stoichiometry of upwelled deep water (Bograd et al. 2015, White et al. 2019) or altered partitioning of organic matter into dissolved or particulate pools under different nutrient scenarios (Carlson et al. 1998). It also does not consider potential systematic changes in the stoichiometry of exported material via each BCP pathway. The small phytoplankton that dominate under nutrient-depleted conditions typically have high C:N and C:P ratios (Baer et al. 2017, Moreno and Martiny 2018). Furthermore, dissolved organic matter and sinking particles both tend to increase in C:N ratio with age (Zakem and Levine 2019). Thus, an alternate hypothesis based on this plug-flow-reactor model might be that increased age before subduction during reduced-wind periods leads to a shift towards communities with high C:N and C:P and increased net CO_2_ uptake. Additionally, it is possible that a plug-flow model is too simplified and that sites of subduction shift further from shore during upwelling periods, thus permitting long surface residence times. These alternate hypotheses make it clear that our ability to predict future change in the BCP is currently limited. Nevertheless, they present predictive frameworks for assessing how changes to physical circulation in EBUSs (which is currently more predictable than biotic responses, Di Lorenzo E and Miller 2017) may alter a key ecosystem service. Testing this prediction, will likely require a coordinated and sustained spatially resolved time series of export flux from the euphotic zone within these biomes.

## Acknowledgments

We are indebted to our numerous colleagues in the CCE-LTER program, and to the captains and crews of the many vessels that have supported the CalCOFI and CCE-LTER Programs. We thank Colleen Durkin, who provided helpful feedback on an early draft. This research was supported by National Science Foundation grants OCE 0548275, OCE-0417616, OCE-1026607, OCE-1637632, and OCE-1614359, and OCE-2224726 to the CCE-LTER Program. Support was also provided by the David & Lucile Packard Foundation.

## Data availability

Data used in this study are available on the CCE LTER DataZoo website: https://oceaninformatics.ucsd.edu/datazoo/catalogs/ccelter/datasets and also deposited at the Environmental Data Initiative: https://edirepository.org/. The CCE LTER data on the Environmental Data Initiative is easily searchable through the CCE LTER landing page at: https://ccelter.ucsd.edu/data/.

## Notes

https://ccelter.ucsd.edu/data/

### Competing Interest Statement

The authors have declared no competing interest.

